# Quantum Algorithm for Metabolic Network Analysis

**DOI:** 10.1101/2025.10.26.684702

**Authors:** Ashish Joshi, Takahiko Koyama

## Abstract

Biological systems, such as cellular metabolism, involve thousands of reactions that together determine how cells grow, respond to their environment, and produce energy. Modeling and analyzing these systems require solving very large mathematical problems that can quickly become computationally prohibitive. To address this challenge, we present a quantum algorithm to analyze metabolic networks, focusing on flux balance analysis as a representative case. We use a quantum interior point method consisting of a quantum subroutine for matrix inversion. Specifically, we reformulate the metabolic optimization problem for efficient execution on a quantum computer using quantum singular value transformation, enabling a rapid solution of complex systems that arise in flux balance analysis. This quantum approach offers a potential computational advantage over classical interior point methods for large and well-conditioned networks. We demonstrate the practical applicability of our method with numerical simulations on the glycolysis and tricarboxylic acid (TCA) cycle network and show that the quantum solution converges to the correct biological objective. This work represents the first application of quantum algorithms to metabolic pathway analysis, establishing a new direction for quantum computational biology and paving the way for quantum approaches to large-scale biological optimization.

## 1 Introduction

Living systems consist of vast networks of interacting molecular processes whose collective behavior determines cellular function and phenotype. Experimental characterization of these systems, while essential, is often time-consuming, expensive, and technically constrained. As a result, mathematical modeling of biological systems has proven to be an important complementary approach. However, unlike idealized physical systems, biological systems are usually not isolated, operate under non-equilibrium conditions, and have numerous interacting components that result in emergent properties that cannot be easily predicted from their individual components. Consequently, considerable effort is being expended to create advanced computational tools to study these systems, including protein folding prediction algorithms like AlphaFold [1], molecular dynamics simulations for drug-target interactions [2, 3], genome-scale metabolic network reconstruction [4], and machine learning approaches for predicting enzyme kinetics and regulatory networks [5]. Together, these advances have enabled applications and translation in personalized medicine [6, 7], drug design and synthetic biology [8, 9], driving breakthroughs in biofuel production [10], antibiotic discovery [11], and optimization of industrial and pharmaceutical manufacturing [12].

Cellular metabolism provides a well-defined example of large-scale biological complexity, comprising interconnected metabolic pathways encompassing all biochemical reactions in an organism [13]. Mathematically, advances have been made in understanding these systems through tools such as flux balance analysis [14], elementary mode analysis [15] and dynamic simulations [16, 17]. While steady state methods like flux balance analysis (FBA) have efficient algorithms that can study genome-scale networks, extending these approaches to larger or more realistic systems, such as multi-species communities or dynamic simulations, remains computationally challenging, underscoring the need for new algorithmic innovations.

In recent years, quantum computing has emerged as an alternative computing paradigm with the potential to exponentially accelerate certain problems that are computationally intractable on a classical computer [18, 19, 20, 21, 22, 23]. Currently, we are in the era of noisy intermediate-scale quantum (NISQ) devices, but with rapid advances in hardware, fault-tolerant quantum computers (FTQC) are expected to become feasible in the coming years [24, 25, 26]. However, similar advances in quantum software are lacking, especially in the field of biology, where even identifying problems that are amenable to a quantum computer is challenging. Metabolic network analysis problems, however, are often constructed from ordinary differential equations, and in the case of FBA, reduces to a linear program [13, 14, 27, 28]. This makes them well-suited for a quantum computing framework, with potential speedup over current classical methods, especially as the problem size grows [29, 30, 31, 32, 33, 34].

In this work, we present the first quantum algorithm specifically designed for FBA of metabolic networks. Given a metabolic network, we use quantum interior point methods (QIPMs) to find the optimal flux [35, 36, 37]. A central part of the QIPM is the solution of a set of linear equations, for which we use quantum singular value transformation (QSVT). Since the runtime of QSVT scales with the condition number of the matrix that is inverted, we use null space projection to reduce this condition number making the algorithm feasible for FBA. With numerical simulations of the carbon cycle, we show that our method converges to the correct optimal solution. As the underlying formulation is directly applicable to general constraint-based metabolic models, the approach naturally extends to genome-scale networks where classical scaling becomes prohibitive. Our work establishes the foundation for quantum-enhanced metabolic network analysis and opens new avenues for studying complex biological systems that are intractable with current computational methods.

## 2 Methods

### 2.1 Flux balance analysis

Metabolic networks describe the complete set of reactions and enzymes involved in cellular metabolism, providing a framework for mathematical analysis and optimization [38, 39, 40, 41]. Here, we focus on flux balance analysis (FBA), which assumes steady-state conditions under which the concentration of metabolites does not change with time [13, 14, 27, 28]. Our goal is to identify the flux vector, the rate of reaction that maximizes a specific biological objective, such as ATP generation or biomass growth. The steady-state assumption reduces the set of ordinary differential equations to a set of linear equations, which can be formulated as a linear programming problem:

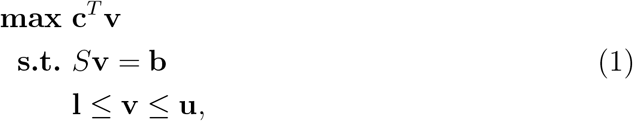

where **v** ∈ ℝ^*n*^ represents the metabolic flux rates, *S* ∈ ℝ^*m×n*^ represents the stoichiometric matrix with mass balance constraints, **c** ∈ ℝ^*n*^ is the biological objective to maximize (for example, biomass production), **b** ∈ ℝ^*m*^ is the rate of metabolite production or consumption (for steady-state assumption **b** = 0) and **l, u** ∈ ℝ^*n*^ are lower and upper bound on the flux rates respectively. We applied the interior point method (IPM) as they are efficient classical algorithms widely used for solving convex optimization problems, such as the linear program problem of FBA [42, 43]. In finding the solution via IPM, first an initial point needs to be found inside the convex polytope of linear constraints. Then, subsequent steps are generated iteratively to converge to the optimal point. This also guarantees that the feasibility condition *S***v** = **b** stays satisfied, as it is satisfied for all points inside the polytope.

We define slack variables to convert the inequality constraints to equality constraints:

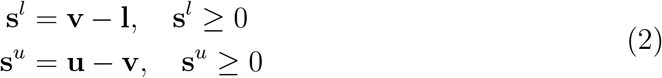

We transform the constrained problem in Eq. (1) to an unconstrained, logarithmic-barrier formulation by adding penalties for each constraint:

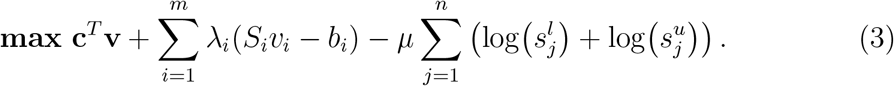

Here, ***λ*** ∈ ℝ^*m*^ are the Lagrange multipliers for the equality constraints. The inequality constraints are transformed into logarithmic barriers and *µ >* 0 is the barrier parameter. As the solution of the linear program lies on the edge of the polytope, *µ* → 0 during optimization. Eq. (3) is also known as the Lagrangian, **ℒ** (**v, *λ***). Any optimal solution of the linear program must satisfy the Karush-Kuhn-Tucker (KKT) conditions [42]:

1. Stationarity:

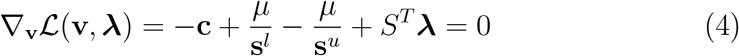
2. Feasiblity:

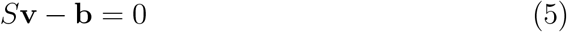

As **ℒ** (**v, *λ***) is strictly convex, these conditions are necessary as well as sufficient.

We use Newton’s method to solve the nonlinear KKT system [42]. To solve a system of equations *F* (**x**) = 0 where *F* : ℝ^*n*^ → ℝ^*n*^ is a vector-valued function, we can expand *F* around a point **x**_*i*_ as

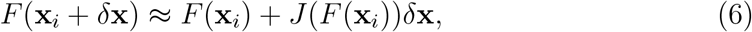

where *J*(*F* (**x**_*i*_)) is the Jacobian matrix. We can solve for *δ***x** by setting Eq. (6) to zero. Thus, the solution can be obtained iteratively using **x**_*i*+1_ = **x**_*i*_ + *δ***x**. For our case, Eq. (4) and Eq. (5) form the set of nonlinear equations

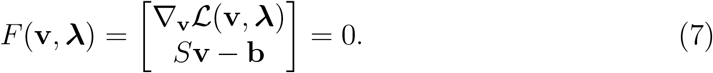

After the Taylor expansion, we get

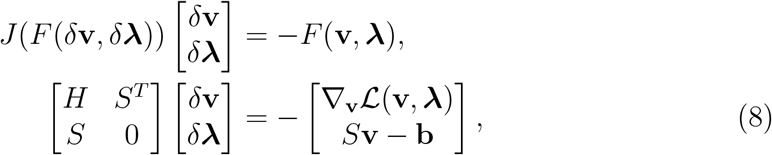

where *H* is the Hessian matrix with only diagonal values 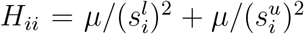. The matrix *J*(*F* (*δ***v**, *δ****λ***)) ∈ ℝ^(*n*+*m*)*×*(*n*+*m*)^ needs to be inverted to solve for *δ***v**. The cost of inverting an *n*-dimensional matrix is 𝒪 (*n*^3^) with the best classical algorithm, and is the most expensive step in solving the Newton’s equation [44]. This can be reduced to 𝒪 (*nsκ*) using the conjugate gradient algorithm for sparse matrices, where *s* is the sparsity and ||*κ* = *A*|||| *A*^−1^|| is the condition number [45]. In the next section we propose a quantum computing routine to improve on this bottleneck.

### 2.2 Quantum interior point method

Quantum interior point methods (QIPMs) aim to solve convex optimization problems using quantum linear solvers (QLSs) [23, 29, 30, 31, 32, 33, 34, 35, 36, 37]. For a square matrix *A* ∈ ℂ^*n×n*^, though extension to any general matrix is straightforward, and a vector **b** ∈ ℂ^*n*^, quantum linear solvers aim to return the solution **x** to the equation *A***x** = **b**. As the vector **b** and matrix *A* are classical data, they need to be encoded into the quantum computer. This is achieved by preparing **b** as a quantum state |**b**⟩ and a unitary operator giving access to the matrix *A*. Then, the quantum linear solver returns a state |**x**⟩ = *A*^−1^ |**b**⟩, where

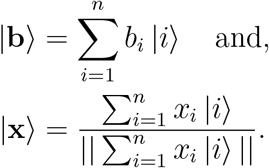

We use quantum singular value transformation (QSVT) based quantum linear solvers [30, 46, 47, 48]. As quantum computers can only perform unitary operations, a general nonunitary matrix needs to be embedded in a unitary matrix. This is obtained via block-encoding [46, 47, 48, 49, 50], in which a unitary operator *U*_*A*_ implements a general matrix *A* by embedding it in a larger unitary matrix such that the upper left block represents *A*, see Fig. 1. After this procedure, we obtain the block-encoding *U*_*A*_ of *A* such that

**Figure 1.**
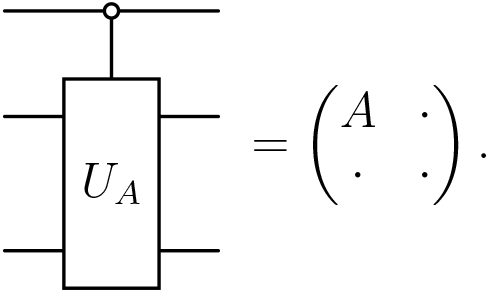
Block-encoding the matrix *A* as a unitary operator in a quantum circuit. *A* is encoded on the top left corner, rest of the blocks can take any values so that *U*_*A*_ is unitary.

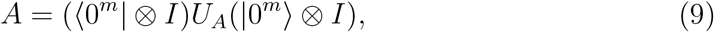

where *m* denotes the number of ancilla qubits required for block-encoding *A*. There are various ways of implementing block-encodings depending on the structure of the matrix *A* [47, 48, 51, 52]. For more details, please see section 7.2.

After block-encoding, the matrix *A* can be applied to a quantum state by applying the unitary *U*_*A*_. However, we want to apply the inverse of the matrix *A* to the vector **b**. This is achieved using QSVT, which applies a polynomial transformation *f* (·) to the singular values of *A*, given its block-encoding *U*_*A*_. By designing a quantum circuit that implements the polynomial transformation as the inverse of the singular values of *A*, we can obtain *A*^−1^. We define *U*_Φ_ as the QSVT circuit that transforms *U*_*A*_ to a block-encoding of *f* (*A*), which is given as

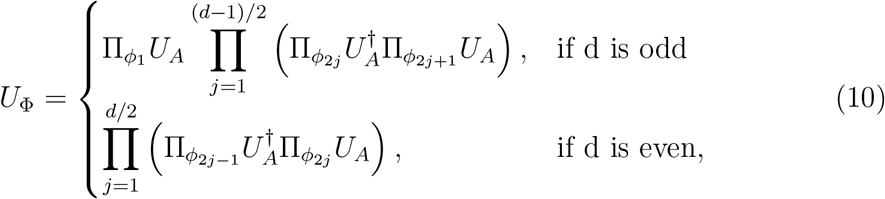

where *d* is the degree of the polynomial *f*, Π_*ϕ*_ = *e*^*iϕZ*^ is the controlled rotation operator acting on the *m* qubits, and Φ = (*ϕ*_1_, *ϕ*_2_, …, *ϕ*_*d*_) is the sequence of angles obtained classically that implements the polynomial transformation to the singular values of *A* [53, 54]. The circuit implementation of QSVT is shown in Fig. 2. For more details on QSVT, please see section 7.3.

**Figure 2.**
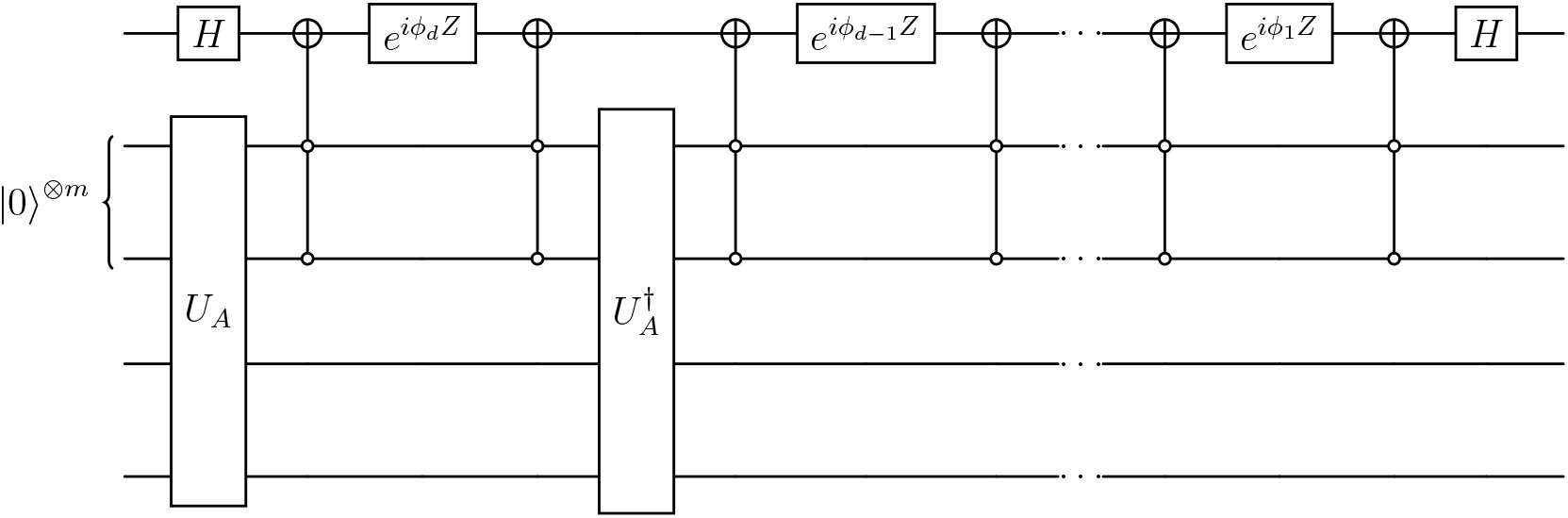
Circuit representing QSVT that implements the desired polynomial transformation *f* (·) on *A* by transforming the block-encoding *U*_*A*_ of A into the block-encoding of *f* (*A*).

In contrast to the classical IPM, QIPM returns a quantum state |**x**⟩ = *A*^−1^ |**b**⟩ as the solution. We performed our numerical experiments with an exact simulator, having full access to the state vector |**x**⟩. For a quantum computer, the use of a quantum tomography routine to extract the vector **x** after each iteration [55, 56] is necessary.

### 2.3 Null space projection

In classical IPMs, the cost of inversion of a general matrix *A* ∈ ℝ^*n×n*^ scales as 𝒪 (*n*^3^). In QIPM, the qubit requirement of block-encoding and QSVT is logarithmic in *n*. However, the query complexity of QSVT scales as 𝒪 (*κ*^2^log(*κ/ϵ*)) [34], where *ϵ* is the desired accuracy and *κ* is the condition number of *A* defined as

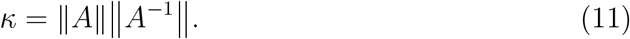

While the dependence of QSVT is only logarithmic in *n, κ* usually increases with *n*. Furthermore, it is known that *κ* → ∞ as the IPM approaches the optimal solution, which could be detrimental for QIPMs [42]. Thus, the scaling of the algorithm depends on the problem-specific parameter *κ*.

Here, we propose a method to reduce *κ* of matrix *A* by adding regularization to its null space projection. Regularization refers to a set of techniques widely used to stabilize numerical algorithms. Since large *κ* is due to a varying orders of eigenvalues in the matrix, one regularization strategy is to add a small diagonal term *λ* to the matrix *A, A* ≈ *A* + *λI*. This procedure, however, does not reduce *κ* to a feasible value for the matrix in Eq. 8 without using a large *λ*, which results in reduced convergence accuracy. We fix this problem by working in the null space of stoichiometric matrix *S* [36, 42]. This results in a smaller but denser projected Hessian matrix, which has a better response to regularization and results in a more stable and accurate algorithm.

For any feasible point **x** *∈ F*, where the feasible set is defined as *F* = {**x** ∈ ℝ^*n*^ : *A***x** = **b**}, the tangent space to the feasible set is the null space of *A*

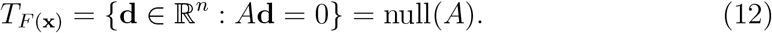

The projector *P* onto the null space of *A* is such that for any **d** ∈ ℝ^*n*^, *P* **d** null(*A*), i.e., we want *P* such that *AP* = 0. Assuming *P* = *I* −*A*^*T*^ *X* for some matrix *X*, we have

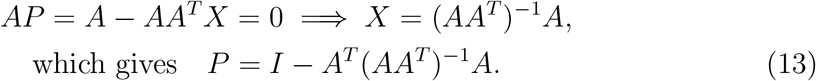

*P* has some nice properties: *P* ^2^ = *P* and *P*^*T*^ = *P*. To apply *P* to the Newton system of equations and replacing *A* with the stoichiometric matrix *S*, we rewrite Eq. (8) as

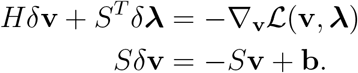

Applying *P* to the upper equation,

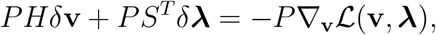

and since *PS*^*T*^ = 0, we obtain

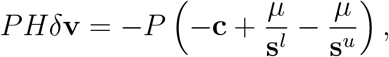

applying *P* again and using *P* ^2^ = *P*, we obtain

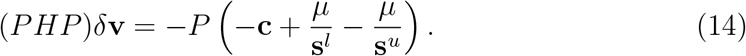

The projected Hessian matrix is regularized as (*PHP* +*λ*_*i*_*I*), where *λ*_*i*_ is the regularization term for *i*th iteration and which depends on the progress of the optimization procedure as well as barrier parameter *µ*_*i*_. For details on the specific implementation, please see section 7.4.

## 3 Results

We tested our method on the carbon core pathway, which consists of two central pathways of cellular energy metabolism, glycolysis and the tricarboxylic acid (TCA) cycle. This network is depicted in Fig. 3, where metabolites are represented in black and enzymes associated with the reaction are represented in red. For this metabolic network, the stoichiometric matrix *S* has dimensions (22 × 20), and we used 6 qubits in our numerically exact simulations (for further details of all the numerical parameters, please refer to section 7.4). Our goal was to find the flux vector that maximizes the production of biomass. The optimization of the objective function Eq. (1) with number of iterations is shown in Fig. 4(a). The black line shows the exact solution obtained classically and the blue line shows convergence of the objective value using QIPM. We also plot the feasibility error, i.e., violation of the equality constraint of Eq. (5), in Fig. 4(b). Thus, the feasibility error stays well within the error margins and we obtain the correct objective value using our approach. Broadly, these results identify the steady-state flux distribution through glycolysis and the TCA cycle that supports maximal cellular growth

**Figure 3.**
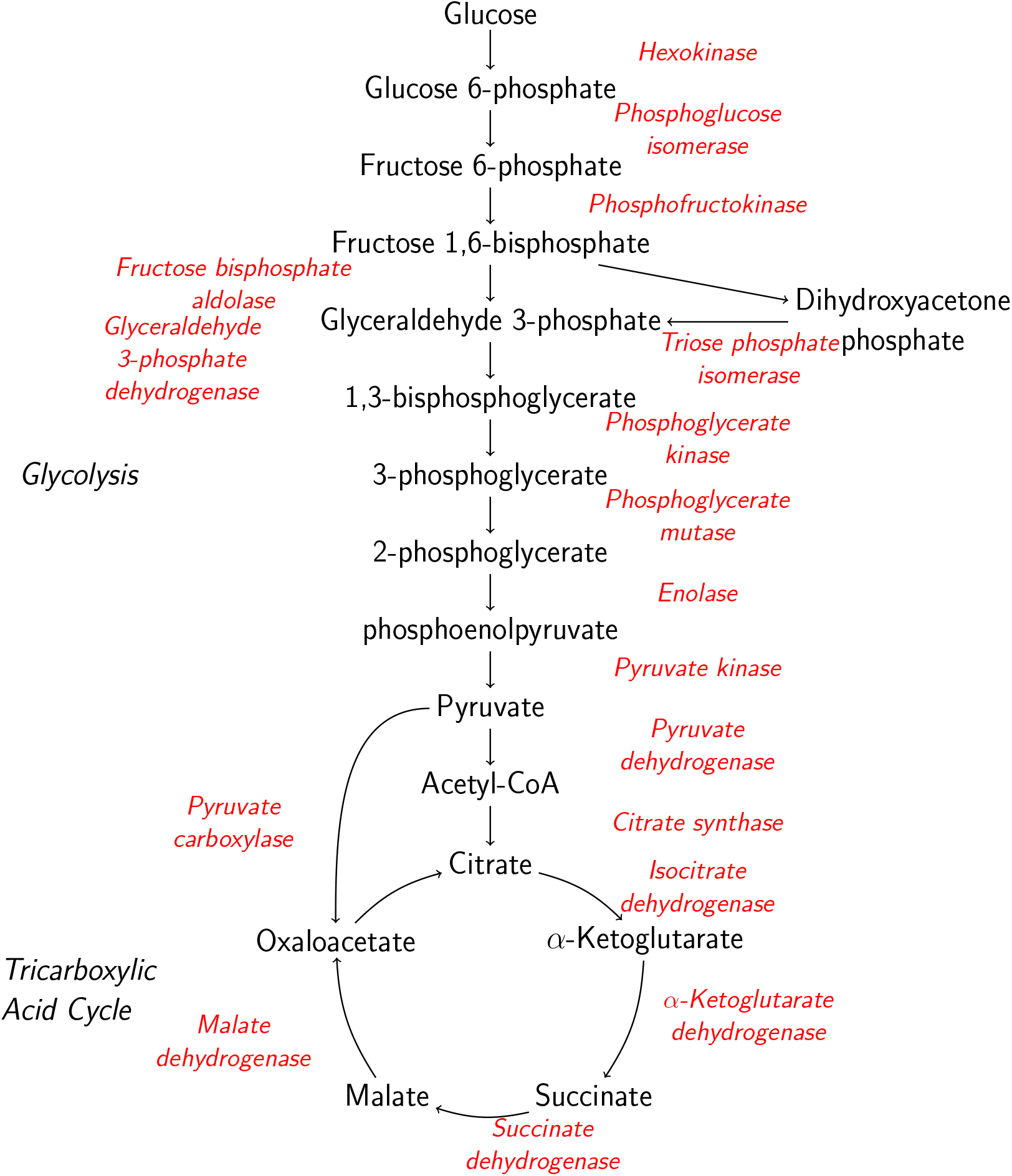
Carbon core pathway used in our numerical experiment, showing the glycolysis and TCA cycle networks. Metabolites are represented in black and enzymes are represented in red.

**Figure 4.**
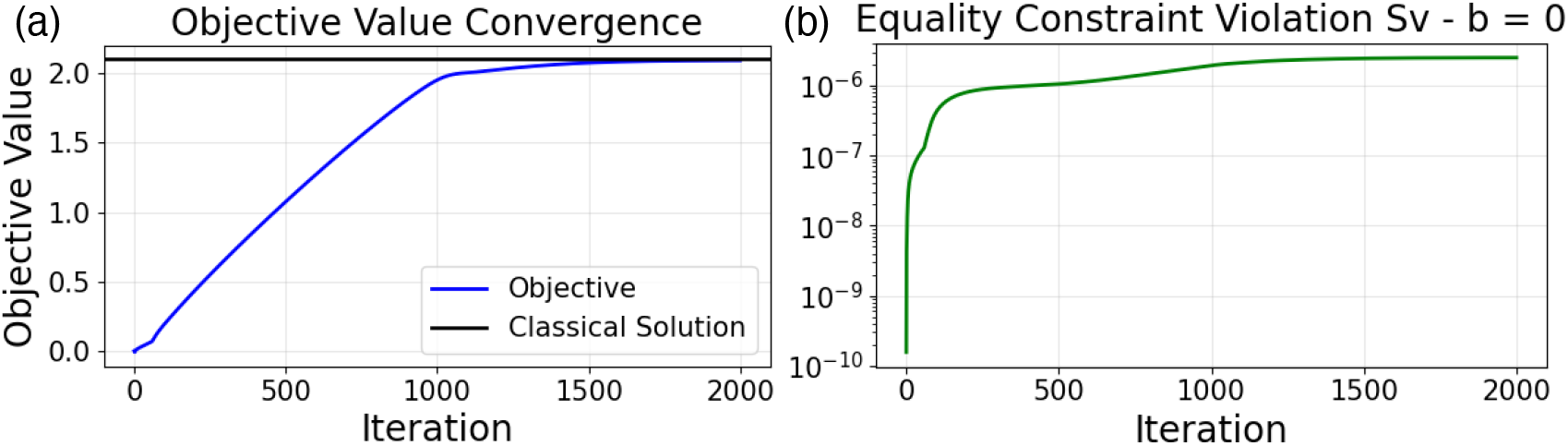
Solution of glycolysis with TCA cycle metabolic network using QIPM. (a) Convergence of the objective value, i.e., maximizing biomass, to the classical solution. (b) Feasibility error, which increases rapidly initially but then stays almost constant.

In Fig. 5(a) we show the routine used for the barrier parameter *µ* and regularization *ϵ*. The objective value reaches the optimal value as *µ* → 0. For numerical stability, it is crucial that the reduction in *µ* with iteration *i* is not severe. Here, we first reduce *µ* slowly as *µ*_*i*+1_ = 0.999*µ*_*i*_ and then as *µ*_*i*+1_ = 0.995*µ*_*i*_. Similarly for regularization *ϵ*, we start from a very high value for numerical stability and then reduce it over the course of optimization to increase accuracy. Akin to machine learning models, *µ* and *ϵ* are hyperparameters and their choice is crucial (see section 7.4).

**Figure 5.**
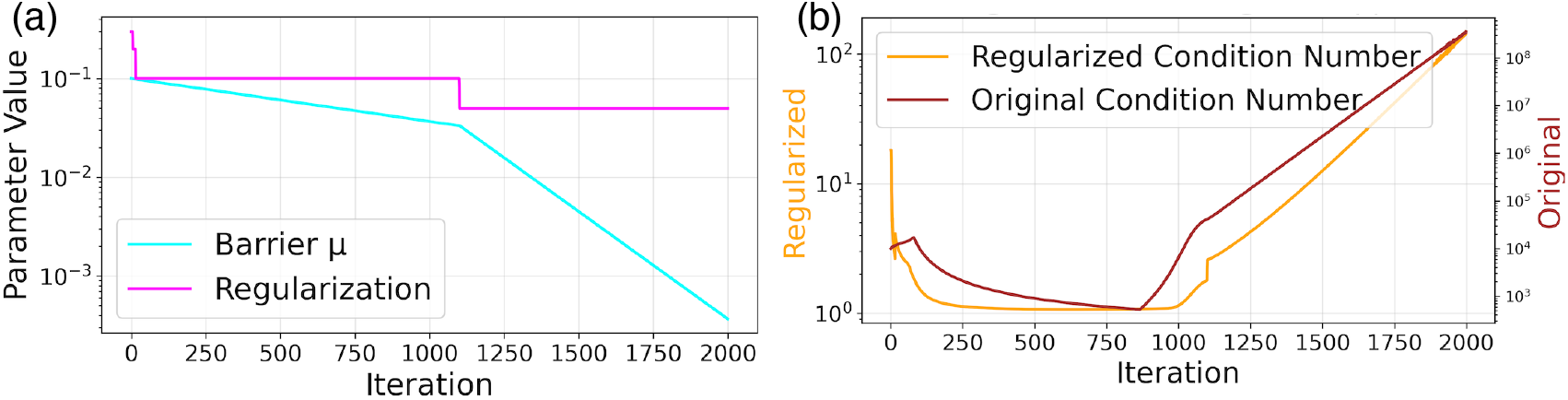
Numerical parameters and condition number during the optimization. (a) Regularization and barrier routine used during the optimization. (b) Condition number of the original matrix and the regularized Hessian.

We show the results of our regularization procedure in Fig. 5(b). The orange plot shows the condition number *κ* of the original matrix in Eq. (8) and the brown plot shows *κ* after the null-space projection procedure and regularization (section 2.3). With this procedure, we see that we could reduce *κ* by many orders of magnitude. We also see that both condition numbers increase rapidly with decreasing *µ*. This might be a problem for larger networks, where *κ* may reach prohibitively large values affecting the accuracy of the matrix inversion via QSVT. By improving numerical stability, this procedure allows the optimization to converge on realistic metabolic flux patterns, ensuring that the predicted reaction rates can be meaningfully interpreted in the context of cellular metabolism.

## 4 Discussion

In this work, we demonstrate the first implementation of quantum interior point methods (QIPMs) to a complex biological system, specifically the optimization of cellular metabolic networks. Using flux balance analysis (FBA), a widely adopted framework for predicting metabolic responses to genetic and environmental changes, we demonstrate how quantum algorithms can identify optimal flux distributions that maximize biologically relevant objectives such as growth or energy yield. Using the stoichiometric matrix representation of glycolysis and the tricarboxylic acid (TCA) cycle, our quantum approach accurately reproduces classical results while laying the groundwork for future studies of larger and more complex networks, where modeling entire-cell or community-scale metabolism remains computationally prohibitive.

As FBA assumes steady-state conditions, it can be solved efficiently by classical algorithms for genome-scale reconstructions of metabolic networks using the classical interior point methods (IPMs) or the simplex method. Thus, the possibility of a quantum advantage for FBA lies mainly in even larger scale networks. A further complication that the current QIPM implementation may face relates to the condition number *κ*, while although problem-specific, does depend on the size of the stoichiometric matrix. This would affect the accuracy of the QSVT matrix inversion, which depends on *κ*. Thus, how well our regularization methods work for large, biologically relevant networks needs to be explored. Furthermore, recent studies have proposed QLSs that improve the dependence on the condition number *κ*, enabling more stable optimization of large-scale metabolic networks [32, 34, 57].

A more realistic representation of metabolism can be achieved by relaxing the steady-state assumption and modeling metabolite concentrations through systems of ordinary differential equations or dynamic FBA. Such models capture regulatory and temporal behavior, for example, how cells adapt their metabolism during growth, stress, or nutrient shifts. This added realism, however, comes at the cost of enormous computational complexity. Classical optimization methods struggle to handle these large, dynamic networks, making them a promising target for quantum acceleration. By demonstrating a quantum approach to metabolic optimization, our work lays the groundwork for modeling metabolism in physiologically realistic contexts, potentially enabling the study of cellular adaptation, metabolic regulation, and community-scale interactions that are currently beyond reach with classical computation. By efficiently exploring high-dimensional biological design spaces, quantum algorithms could help identify alternative flux pathways and model complex metabolic interactions within microbial communities, ultimately enabling personalized interventions such as microbiome optimization and early detection of disease through metabolic biomarkers

## 5 Conclusion

In this study, we have demonstrated a quantum algorithm for metabolic pathway analysis that can be implemented on early fault-tolerant quantum computing (FTQC) devices with modest qubit counts and circuit depths. Although we employed flux balance analysis (FBA) for the current demonstration, our approach is extendable beyond this framework. More realistic simulations of multi-species communities will require dynamic models utilizing systems of differential equations to capture continuous metabolite exchanges as the system evolves toward equilibrium. As quantum computers advance in qubit count, circuit depth, and gate fidelity, these mechanistic biological models can be simulated at runtimes unattainable with classical methods. This work represents one of the first biologically relevant quantum algorithms, designed specifically for near-term FTQC implementation, establishing practical underpinnings for quantum computational biology as fault-tolerant hardware emerges.

## 6 Acknowledgements

Human Biology-Microbiome-Quantum Research Center (Bio2Q) is supported by World Premier International Research Center Initiative (WPI), MEXT, Japan. This work was also supported by the Center of Innovation for Sustainable Quantum AI (JST Grant Number JPMJPF2221). The authors are grateful to Scott Behie for valuable discussions and comments on the manuscript.

## 7 Appendix

## 7.1 Quantum computing background

As this work is intended for readers from diverse backgrounds, in this section we briefly describe the quantum physics concepts and notations used throughout the text. For more information, we direct the readers to detailed references on quantum computing [18, 23].

The state of quantum system is denoted by the wave function |*ψ*⟩. For a single qubit, the quantum state can be written as a column vector in the complex vector space ℂ^2^, i.e., with two dimensions, and can be represented as

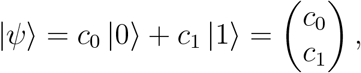

where |0⟩ and |1⟩ are the orthonormal basis states and *c*_0_, *c*_1_ *∈* ℂ are known as probability amplitudes. The state |ψ⟩ is in a superposition of the two basis states and upon measurement will result in the state |0⟩ with a probability ∥*c*_0_∥^2^ and the state |1⟩ with a probability ∥*c*_1_∥^2^. For *n* qubits, the quantum state lives in a 2^*n*^-dimensional complex vector space and can be represented as

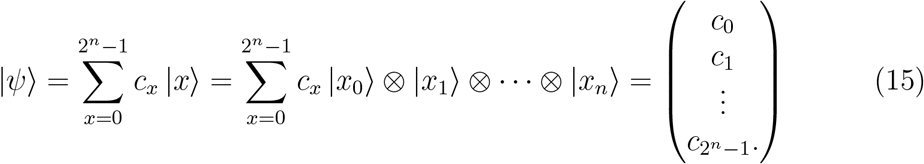

The orthonormal basis |0⟩, |1⟩, …, |2^*n*^ − 1⟩ is called the computational basis. The Hermitian adjoint of the state |*ψ*⟩ is given by 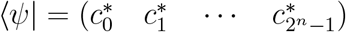 which is a row vector.The inner product between two quantum states |*ψ*⟩ = ∑ _*x*_*a*_*x*_|*x*⟩ and |*ϕ*⟩ = ∑ _*x*_*b*_*x*_|*x*⟩ is given by

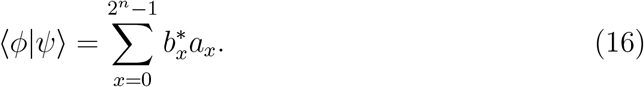

The inner product of a state with itself is the norm of the quantum state

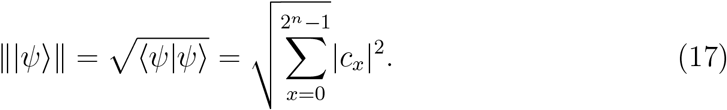

Physical observables are represented as operators, which are linear transformations corresponding to 2^*n*^ × 2^*n*^ matrices for *n*-qubits. An observable *O* can be written as

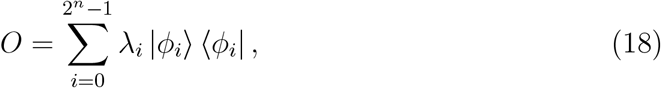

where the eigenvalues *λ*_*i*_ are real if *O* is Hermitian and the eigenvectors |*ϕ*_*i*_⟩ form an orthonormal set. In a quantum system, we only have access to the observables, usually through the expectation value

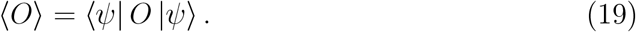

In quantum computing, we work with quantum circuits and quantum gates, which are analogues to the classical Boolean circuits and gates. Quantum gates are unitary operators and allow us to manipulate quantum states. Some essential gates are:

- The Pauli gates *X, Y, Z* and the identity gate *I*

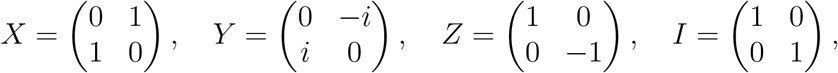
- The Hadamard gate *H* and the *T* gate

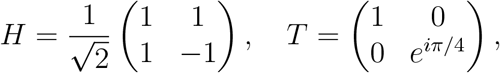
- The CNOT gate

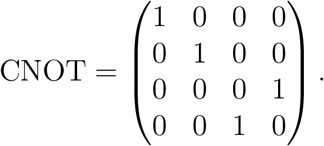

An n-qubit gate is a 2^*n*^ × 2^*n*^ matrix, given by the tensor product of individual single-qubit gates. The quantum circuit then consists of initializing a quantum state, typically to a product state like |0⟩ ^⊗*n*^, applying a sequence of quantum gates and measuring the qubits in the computational basis.

## 7.2 Block-encoding

Given a general square matrix *A* ∈ ℂ^*n×n*^, the unitary operator *U*_*A*_ is an (*α, m, ϵ*)-block-encoding of *A* if

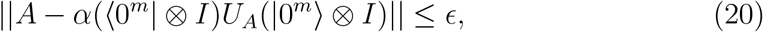

where *α* is the normalization constant for *A, m* is the number of ancilla qubits used for block-encoding *A* and *ϵ >* 0 is the error. The matrix *A* can now be applied to a quantum state by applying *U*_*A*_. Circuit implementation of block-encoding depends on the structure of matrix A, please refer [23] for a detailed review of the various methods. Here we mention some common cases.

- The simplest case is of a unitary matrix, which is a (1, 0, 0)-block-encoding of itself.
- The sparse access model defines how to create block-encodings for sparse matrices. For a matrix *A* ∈ ℂ^*n×n*^ with *s*_r_ -row sparsity and *s*_c_ -column sparsity and ∥*A*∥_max_ = max_*i,j*_|*A*|_*ij*_, 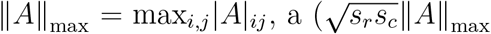, *n* + 3, *ϵ*)-block-encoding of *A* can be defined using oracles *O*_*r*_, *O*_*c*_, and *O*_*A*_ as

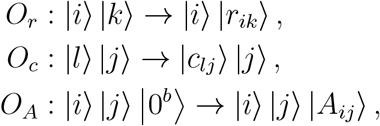

where *i, j ∈* [2^*n*^] − 1, *k ∈* [*s*_*r*_], *l ∈ s*_*c*_, *r*_*ij*_(*c*_*ij*_) is the index of the *j*-th (*i*-th) nonzero entry in the *i*-th (*j*-th) row (column) of *A* and |*A*_*ij*_⟩ is a *b*-bit encoding of *A*_*ij*_ [23, 48].
- For a Gram matrix *A* with *A*_*ij*_ = ⟨*ψ*_*i*_ |*ϕ*_*j*_⟩, a (1, *m*, 0)-block-encoding can be defined as

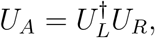

where *U*_*L*_ and *U*_*R*_ are state preparation unitaries. This method can be extended to general matrices [23, 48].

## 7.3 Quantum singular value transformation

We begin with a brief description of quantum signal processing (QSP), which builds up to the QSVT [48]. Given a signal rotation operator *W* and a signal-processing rotation operator *S*,

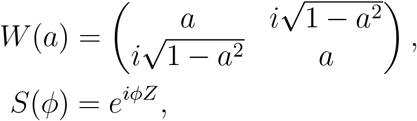

The QSP sequence is defined as

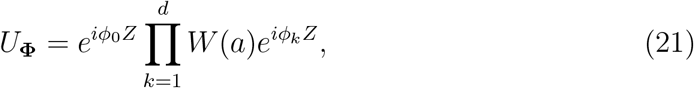

where **Φ** = {*ϕ*_0_, *ϕ*_1_, …, *ϕ*_*d*_} is a tuple of phases. Then there exists a ***ϕ*** which can implement a polynomial transformation to the signal *W* (*a*) as

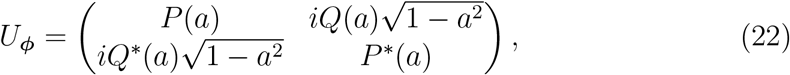

for *a ∈* [−1, 1] such that:

1. *deg*(*P*) ≤ *d, deg*(*Q*) ≤ *d* − 1,
2. *P* has parity *d* mod 2 and *Q* has parity (*d* − 1) mod 2,
3. |*P*|^2^ + (1 − *a*^2^)|*Q*|^2^ = 1.

By defining a 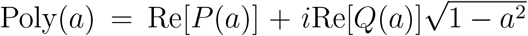 and with some mild constraints on Poly(*a*), we can accurately approximate any real polynomial by selecting proper *P* and *Q*.

QSVT generalizes QSP from a single-qubit operation to general matrices. Given a square matrix *A*, its singular value decomposition (SVD) can be written as

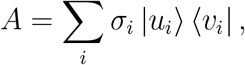

where *σ*_*i*_ are the singular values of *A*. Given a (1, *m*, 0)-block-encoding *U*_*A*_ of *A* such that

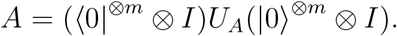

We define the projector operators that can project *A* from *U*_*A*_ as Π =∑_*i*_ |*v*_*i*_⟩ ⟨*v*_*i*_| and 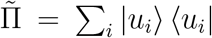 such that 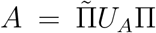. Then, we can apply a polynomial transformation Poly(*A*) to the singular values of *A* using *U*_**Φ**_, where

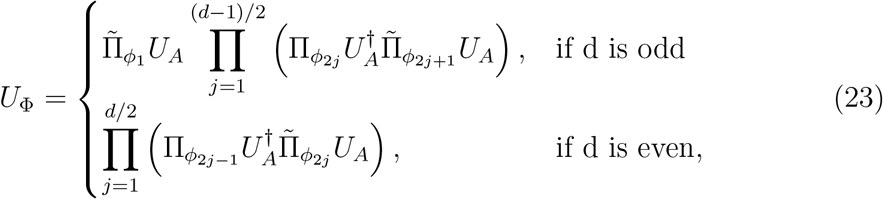

such that

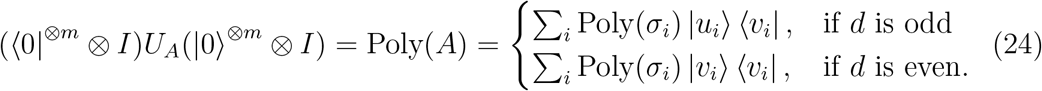

For details on the full derivation, please refer to [30, 48].

## 7.4 Parameters for numerical experiments

The proposed QIPM algorithm can be divided in two parts: an outer part of the algorithm that is classical and is similar to classical IPMs and an inner part that is matrix inversion using QSVT. The outer classical loop defines all the parameters used in our numerical experiments. The problem of finding a feasible initial point is much easier than finding the solution of the linear program, and we obtain this initial point classically [58]. We start with the barrier parameter *µ* = 0.1, which reduces with every iteration *i* as *µ*_*i*+1_ = 0.999*µ*_*i*_ for roughly the first half of the optimization procedure and as *µ*_*i*+1_ = 0.995*µ*_*i*_ for the rest of the procedure. If during the optimization the solution vector gets too close to the boundary of the polytope described by linear equations, the condition number *κ* can increase rapidly. To prevent this and maintain numerical stability, we added a minimum slack value *s*_min_ = *µ*, so that 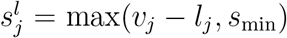 and 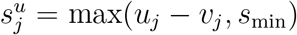. The optimal solution lies on the boundary at one of the corners and with *µ* → 0 with number of iterations, we also have *s*_min_ → 0.

For our purposes the most important property of the matrix to be inverted is its condition number *κ*. As mentioned in the main text, we use the null space projection method to keep *κ* low and in exchange we end up with a denser matrix. To further improve *κ*, we add a small term *λ* to the diagonal entries of the matrix as regularization. Larger regularization leads to a more stable but less accurate algorithm. We start with a large regularization to ensure numerical stability and then reduce it over the course of the optimization (Fig. 5(a)). The final parameter for the outer loop is the step size *α*, which is used to update the solution vector at each iteration **v**_*i*+1_ = **v**_*i*_ + *αδ***v**. We set *α* to change with the regularization *λ*, so that *α* = 0.2 for *λ >* 0.1, *α* = 0.7 for *λ* = 0.1 and *α* = 1.0 for *λ <* 0.1.

We performed our numerical experiments using Pennylane’s default qubit simulator, which is an exact state vector simulator [59]. For the carbon core pathway, the stoichiometric matrix has dimensions (22 × 20), the Jacobian (Eq. (8)) has dimensions (42 × 42) and the null space projection creates a matrix of dimensions (22 × 22) to be inverted. For this matrix we used 6 qubits and performed block-encoding exactly. The angles to implement the polynomial approximation were generated using pyqsp [30, 60, 61]. The degree of polynomial required to approximate 1*/x* depends on *κ* and is not known beforehand. Here, for the purpose of tracking *κ* over the course of optimization, we calculate it explicitly.

